# Mussel memory: bivalves learn to fear parasites

**DOI:** 10.1101/2021.08.28.456938

**Authors:** Christian Selbach, Loïc Marchant, Kim N. Mouritsen

## Abstract

Fear plays a crucial role in predator-prey interactions and can have cascading impacts on the structure of whole ecosystems. Comparable fear effects have recently been described for hosts and their parasites but our understanding of the underlying mechanisms remains limited by the lack of empirical examples. Here, we experimentally tested if bivalves *Mytilus edulis* can ‘learn to fear’ the infective transmission stages (cercariae) of the trematode *Himasthla elongata* and if experienced mussels change their parasite-avoidance behaviour accordingly. Our results show that previous experience with parasites, but not infection per se, lead to a reduced filtration activity in mussels in the presence of cercariae compared to parasite-naïve conspecifics. This reduction in filtration activity resulted in lower infection rates in mussels. Since parasite avoidance comes at the cost of lower feeding rates, mussels likely benefit from the ability to adjust their defence behaviour when infection risks are high. Overall, these dynamic learning processes of avoidance behaviour can be expected to play a significant role in regulating the bivalves’ ecosystem engineering function in coastal habitats.

## Introduction

Fear saves lives, and fear shapes ecosystems. Fear can be defined as a central state that is induced when an organism perceives a threat or danger, and triggers physical or behavioural responses (Adolphs 2013; Silva et al. 2016). Although such behavioural or physical responses are energetically costly, they increase fitness and have emerged as evolutionary stable strategies in different taxa (Brown et al. 1999). For example, a herd of elk maintains a constant vigilance against wolves, or a school of fish moves in complex collective manoeuvres to confuse and avoid potential predators (Laundré et al. 2010; Marras & Domenici 2013). Such fear-induced predator avoidance behaviours can lead to non-consumptive or trait-meditated indirect effects on other organisms in a community, and can have cascading impacts on the structure of whole ecosystems, comparable to or even exceeding the direct effects of predation (Peacor & Werner 2001; Ripple & Beschta 2004; Preisser et al.; 2005; Creel & Christianson 2008; Suraci et al. 2016). Predator avoidance behaviour of herbivores (antelopes) in the African savannah, for instance, has been shown to shape the distribution and structure of Acacia tree communities in these habitats (Ford et al. 2014). These ecological consequences of predator avoidance are referred to as the ‘ecology of fear’ (Brown et al. 1999; Zanette & Clinchy 2019).

Similar to predators, parasites can induce ‘fear’ or ‘disgust’ in their hosts (Buck et al. 2018; Weinstein et al. 2018). Free-living organisms have developed a variety of defence and avoidance strategies and mechanisms against parasites or pathogens, ranging from evading parasite transmission stages to avoiding interaction with infected conspecifics, or staying clear of risky infection hot-spots, such as areas contaminated with faeces (Mouritsen 2017; Behringer et al. 2018; Buck et al. 2018; Weinstein et al. 2018; Daversa et al. 2021). The ecological impacts of this fear of parasites have been shown to be comparable to the cascading effects of predator avoidance (Rohr et al. 2009). For instance, larval amphibians increase their activity to escape infective trematode cercariae in the water. This in turn not only impairs the tadpoles’ grazing activity and growth rates, but also their susceptibility to predation, and the resource availability for other grazers, thereby affecting the food web dynamics in the ecosystem (Marino & Werner 2013; Buck et al. 2018).

Overall, the capacity to avoid risks from predators or parasites requires that organisms have an intrinsic ability and/or can learn to differentiate between dangerous and safe conditions (Laundré et al. 2010). So far, most of the evidence that host organisms can learn to fear and avoid their parasites comes from terrestrial insects or mammals, while in aquatic systems, risk learning has been almost exclusively studied for predator-prey interactions (Behringer et al. 2018 and reference therein). Results suggesting avoidance learning of parasite threats in aquatic environments come from individual studies on trematode-fish interactions (James et al. 2008; Klemme & Karvonen 2016). Accordingly, our understanding of these fundamental interspecific interactions in aquatic habitats is currently limited by the lack of empirical examples from most host-parasite systems. Changes in parasite avoidance behaviour as a result of experience and learning can have significant implications for disease dynamics and central ecological functions, and need to be explored for further ecologically important host-parasite systems in aquatic environments (Behringer et al. 2018; Selbach & Mouritsen 2020).

Here, we use the bivalve *Mytilus edulis* and its trematode parasite *Himasthla elongata* as a model system to study the capability of hosts to learn to avoid parasites, and assess the ecological implications of these processes. Blue mussels *M. edulis* are an important keystone species in Atlantic intertidal communities. They form extensive mussel beds along the coastline where they fulfil central ecological roles, such as filtering out organic matter and creating biogenic reefs, on which other organisms depend for shelter, substrate and foraging (Ragnarsson & Raffaelli 1999; Bertness 2007; Commito et al. 2008; Larsson et al. 2017). The trematode *H. elongata* utilizes blue mussels as an intermediate host. Mussels become infected via free-swimming parasite dispersal stages, the cercariae, that are emitted from the common periwinkle snail *Littorina littorea* (Werding 1969). Once inside the mussel, the parasites encyst as metacercariae in the host tissue and can reduces growth rates (Bakhmet et al. 2017), and render the mussel more vulnerable to predation (Lauckner 1983). Recent findings have shown that blue mussels try to avoid parasite infections by rapidly contracting their siphons and closing their shells when sensing cercariae in the water column (Selbach & Mouritsen 2020). However, this parasite avoidance strategy comes at a cost to the mussels, as it prevents them from filtrating and feeding during the parasite-avoidance phase (Mouritsen et al. 2021).

In a set of microcosm experiments, we tested if mussels can learn to fear infective parasite stages in the water, and if this learning process can modulate behavioural changes in their filtration activity in the presence of parasites. We hypothesised that mussels with previous parasite encounters would ‘learn to fear’ this threat and exhibit stronger avoidance behaviours compared to naïve conspecifics. Overall, this could have important ecological implications, e.g., as it would impact the susceptibility of naïve vs experienced mussels to parasite infections and thereby affect the distribution of parasites in an ecosystem. Moreover, it could have important implications for the ecosystem engineering potential of blue mussels, as mussels that gradually learn to avoid parasites by shutting down their filtration activity would remove less organic matter from the environment when parasites are present.

## Material and methods

### Collection of samples

Blue mussels *Mytilus edulis* were provided by the Danish Shellfish Centre, Mors, Denmark. These mussels come from deeper sublittoral waters of the Limfjord, Denmark (56°53’29.2”N 9°09’58.0”E) where no *Littorina littorea* occur, ensuring that no mussels were infected by *Himasthla elongata* (Buck et al. 2005). A subsample of mussels was dissected to confirm the absence of trematode infections. All mussels were brought to the Marine Biological Station, Rønbjerg harbour, Limfjorden, Denmark, where all further treatment and experimentation was carried out. To obtain the parasites for infection experiments, periwinkle snails *L. littorea* were collected at an intertidal zone near Knebel in eastern Jutland, Denmark (56°12’32.2”N 10°28’47.2”E). Snails were screened for patent infections with *H. elongata*, and infected snails were isolated and kept dry in a climate chamber at 16°C until the start of the experiment.

### Experimental infection of mussels

To obtain infected *M. edulis* for the treatment groups, we experimentally infected mussels with freshly emitted cercariae of *H. elongata*. For this, 80 mussels were established in a 20L aquarium with running sea water and air supply. Infected periwinkles were individually placed in glass jars with sea water at approximatively 25°C and placed under a light source to induce cercarial shedding. After 2 h, the glass jars were examined for released cercariae, and those containing free-swimming *H. elongata* were emptied into the aquarium containing mussels. To allow a gradual build-up of trematode infections, mussels were repeatedly exposed to cercariae over the course of several days. The last batch of parasites was administered 24 h before the experiment. A second batch of mussels (n=80) was treated the same way but received only filtered sea water instead of parasites (naïve mussels, see below).

To obtain cercariae for the exposure treatments (see below), infected *L. littorea* were placed in glass jars with sea water, as described above. After 2 h, *H. elongata* cercariae were moved to a petri dish and separated in groups of 200 cercariae using a glass pipette. Each batch of cercariae was kept in 10 ml of sea water in a small petri dish. Cercarial shedding and counting was done within 2 h before the start of the experiment, ensuring all cercariae were ≤2 h old and fully active.

### Experimental design

To test if blue mussels can learn to change their parasite-avoidance behaviour based on previous experience, we performed filtration experiments with pre-infected and naïve mussels (i.e., without previous experience of parasites), either in the presence (‘exposed’ treatment) or absence (‘unexposed’) of trematode cercariae. Each treatment consisted of 10 replicates, resulting in a total of 40 samples. The experiment was carried out in a climate chamber at 16°C and performed in two batches of 20 mussels, five per treatment, over two days. To avoid confounding effects of mussel size, only individuals of similar size were used in the experiment (mean ± SD = 34.3 ± 1.5 mm, n=40).

Mussels were individually placed in a 1.5L bucket with 1.1L of filtered sea water and an air stone for oxygenation and allowed to acclimatize for 2 h before the start of the experiment. After the acclimation period, 0.4L of water containing approximately 30 million cells of algae (TETRASELMIS 3600, Instant Algae, Reed Mariculture) was added to each bucket, resulting in a final concentration of 20 million cells per litre. The exposure treatments each received 10 ml of sea water containing 200 *H. elongata* cercariae; the non-exposure treatments received the same amount of filtered sea water without cercariae. Two additional buckets per day received sea water and algae concentration without the addition of mussels or cercariae to account for any possible sedimentation or breakdown of algae cells during the course of the experiment.

After 2 h, all mussels were removed from the buckets and, after careful stirring, 0.5L of water was removed from each bucket. Water samples were vacuum filtered through a 0.45 µm GC50 Glass Fibre Filter (Advantec, Japan) and filters were immediately frozen at -18°C for later chlorophyll-a analyses. Mussels were individually placed in a small bucket with filtered sea water for 12 h to allow metacercariae to establish before being frozen at -20°C until dissection.

### Parasite infection intensity

Mussels were measured, dissected and the soft tissue tightly squeezed between two glass plates to quantify the infection intensity of *H. elongata*. Metacercariae were counted in the different mussel tissues under a stereomicroscope (ZEISS Stemi 2000c, Germany).

### Measurements of chlorophyll-a concentration

Filters containing retained microalgae were transferred to dark test tubes together with 5 ml 96% ethanol and stored dark for 18 h at room temperature. The samples were then placed in an ultrasonic cleaner for 5 min followed by 20 sec on a minishaker and finally 10 min centrifugation (4.000 rpm). The resulting supernatant of each sample (3 ml) was transferred to a spectrophotometer (Spectronic Helios Alpha®) and the absorbance was measured at 750 and 665 nm. Subsequently, the chlorophyll-a concentration (μg L^-1^) was calculated according to Riemann (1989).

### Data analyses

Statistical analyses were carried out in Statistical Package for the Social Science (IBM SPSS 27.0). All analyses were preceded by tests of homogeneity of error variance (Levene’s test) and evaluation of normality. Three replica (i.e., individual mussels) were excluded in the final analyses: two mussels spawned heavily during the experiment, potentially affecting their filtration activity, and one mussel was excluded due to unreliable spectrophotometric absorbance measurement (at 750 nm). These omissions resulted in effective sample sizes of 9-10 per treatment.

Preliminary full model 3-way ANOVA, entering chlorophyll-a concentration as dependent variable and parasite-exposure (absence/presence of cercariae), infection status (pre-infected/uninfected) and experimental day as fixed factors, demonstrated neither day-interaction nor overall day-effect (F_1,29_ ≤ 0.758, P ≥ 0.391). Hence, data from the two experimental days were pooled in subsequent analyses in order to optimize statistical power. To test for differences between treatments, a full factorial 2-way ANOVA was performed with post-experimental chlorophyll-a concentrations as dependent variable and parasite-exposure and infection status as fixed factors.

## Results

The 2-way ANOVA, entering chlorophyll-a concentrations as dependent variable and parasite-exposure and infection status as fixed factors, showed significant two-way interaction as well as main effects (Table 1, Fig. 1). Overall, parasite exposure caused a substantial reduction in the mussels’ filtration activity (i.e. high chlorophyll-a concentration post-experimentally) explaining 41.5% of the variance (Table 1). This effect was particularly pronounced among pre-infected mussels that left a 46% higher chlorophyll-a concentration than parasite naïve mussels (Fig. 1). Infection per se had no bearing on the mussels’ filtration, as evident from the identical chlorophyll-a concentrations for unexposed parasite naïve and unexposed pre-infected mussels (Fig. 1). For the group of parasite-naïve mussels, those exposed to *Himasthla* cercariae during the experiment left a 33% higher chlorophyll-a concentration than unexposed individuals. This difference was non-significant though (Fig. 1).

**Table 1.** Summary statistics of full model 2-way ANOVA including post-experimental chlorophyll-a concentration (μg L^-1^) as dependent variable and parasite exposure (presence/absence of *Himasthla elongata* cercariae) and infection status (pre-infected/parasite naïve) as fixed factors. Partial η^2^ denotes effect size, i.e. the proportion of variance explained.

| Source | df | F | P | Partial $\eta^2$ |
| --- | --- | --- | --- | --- |
| Parasite exposure | 1 | 23.365 | <0.0005 | 0.415 |
| Infection status | 1 | 6.338 | 0.017 | 0.161 |
| Interaction | 1 | 5.139 | 0.030 | 0.135 |
| Error | 33 |  |  |  |

**Figure 1.**
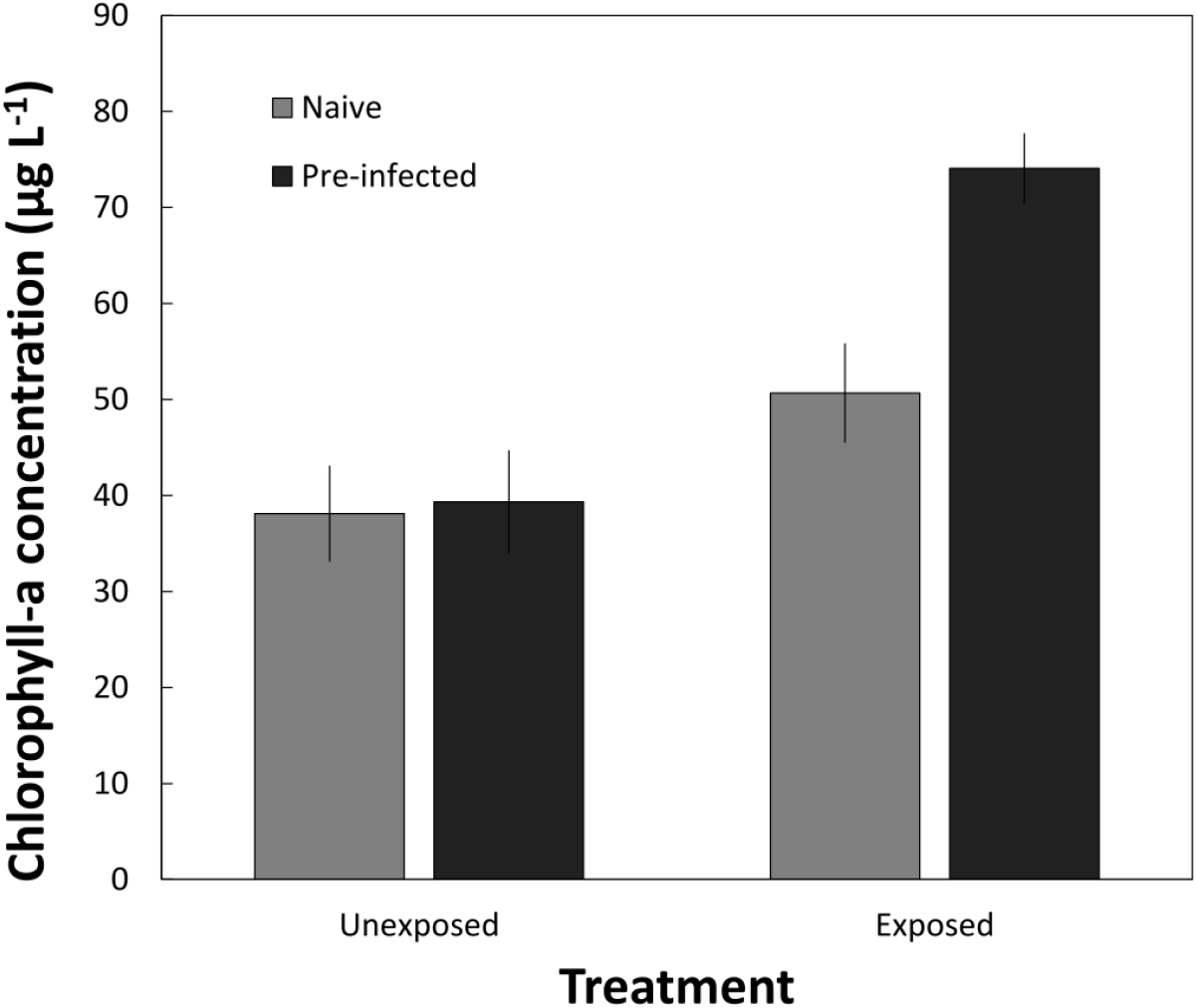
Post-experimental chlorophyll-a concentration (mean μg L^-1^ ± SE) after 2 hrs of filtration by blue mussels *Mytilus edulis* exposed and unexposed to *Himasthla elongata* cercariae. Naïve: pre-experimentally uninfected mussels; Pre-infected: mussels infected by *H. elongata* prior to the experiment. N = 9-10 for each treatment combination. See Table 1 for summary statistics (main analysis). Turkey HSD post-hoc tests: unexposed, naïve versus pre-infected (P = 1.000); exposed, naïve versus pre-infected (P = 0.009); parasite naïve, unexposed versus exposed (P = 0.333); pre-infected, unexposed versus exposed (P < 0.0005).

Focusing on parasite-naïve mussels experimentally exposed to infective *Himasthla* larvae, there was a significant negative linear relationship between the post-experimental parasite load (log-transformed) and chlorophyll-a concentration (Fig. 2). This demonstrates that the rate by which parasite-exposed mussels acquired infections increases exponentially with filtration activity.

**Figure 2.**
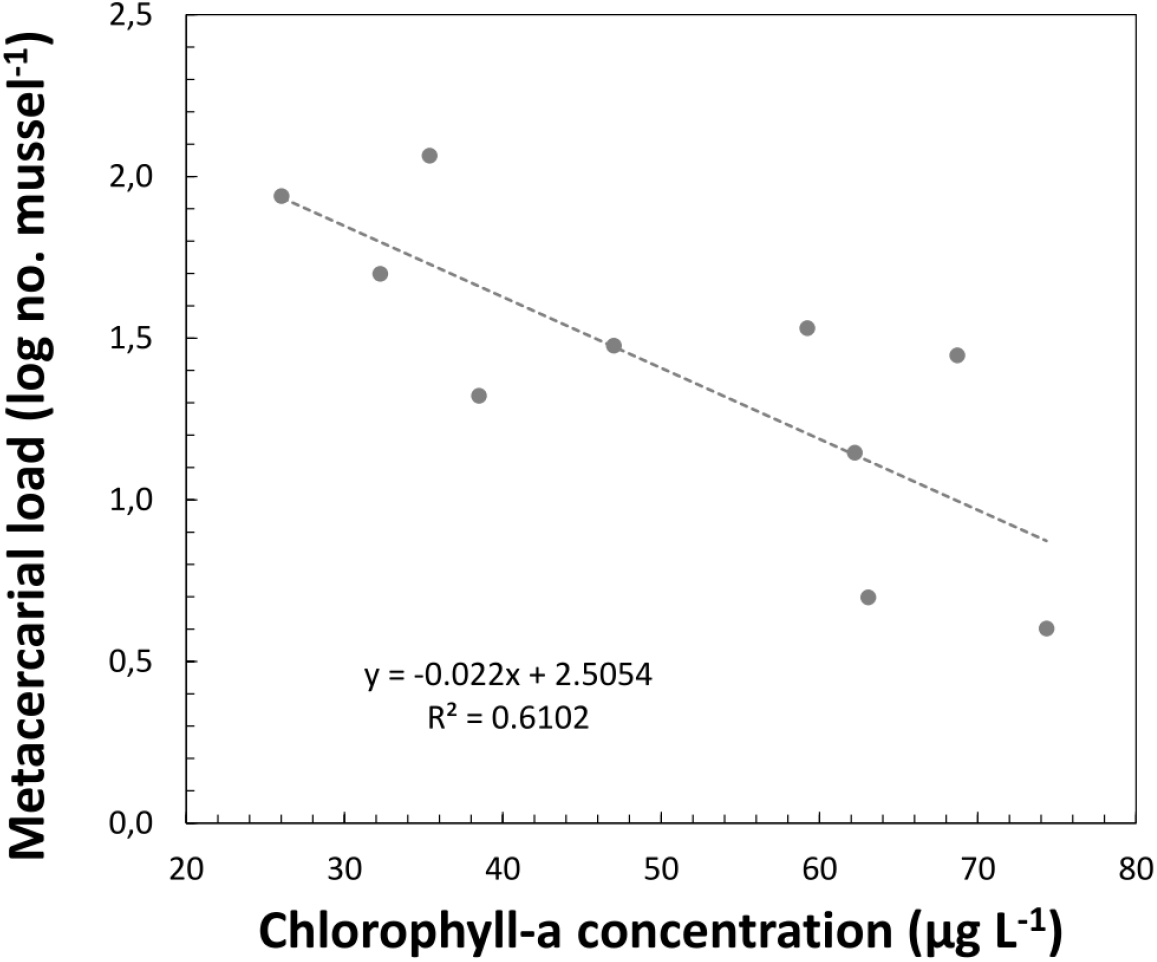
Relationship between post-experimental total metacercarial load in blue mussels *Mytilus edulis* (log-transformed no. ind.^-1^) and chlorophyll-a concentration (μg L^-1^) for pre-experimentally uninfected (naïve) mussels exposed to *Himasthla elongata* cercariae. Linear regression: r^2^_8_ = 0.610, P = 0.008).

## Discussion

Associative learning and risk avoidance have been shown for a range of predator-prey interactions but have received considerably less interest for host-parasite systems (Klemme & Karvonen 2016). Accordingly, our understanding of host learning in parasite avoidance remains limited by the lack of empirical examples. In this study, we tested if blue mussels *M. edulis* can ‘learn to fear’ the infective cercariae of the trematode parasite *H. elongata* in the environment, and if experienced mussels change their parasite-avoidance behaviour accordingly. Our results show that previous parasite encounters, but not established trematode infections in the host, lead to a reduced filtration activity in mussels in the presence of cercariae compared to parasite-naïve conspecifics. This confirms our hypothesis that bivalves can ‘learn to fear’ their parasites, and consequently adjust their behaviour to avoid additional infections. The reduced infection rates in mussels that showed lower filtration activities demonstrate the effectiveness of this avoidance reaction. The absence of any difference in chlorophyll concentration between infected and naïve mussels in the unexposed treatment show that established infections of *H. elongata*, which predominantly encyst in the mantle and foot tissues (Lauckner 1983), do not directly impact the mussels’ filtration ability. Non-consumptive effects of *H. elongata* on *M. edulis* and the reduction of the mussels’ filtration activity as a result of parasite-induced avoidance behaviour have recently been discussed (Selbach & Mouritsen 2020, Mouritsen et al. 2021). The results of our study now highlight the dynamic nature and plasticity of this interaction.

A parasite avoidance strategy that comes at the price of reduced filtration activity and feeding time can be considered highly costly for mussels. For instance, reduced energy uptake due to defensive anti-parasite responses will likely result in lower growth rates and higher risks of predation, since smaller bivalves are more accessible to a wider range of predators (Mascaró & Seed 2001; Larsen et al. 2013). A similar trade-off between parasite avoidance and other threats (predation and competition) could be demonstrated for amphibian tadpoles (Koprivnikar & Penalva 2015; Buck et al. 2018). Overall, such parasite-defence strategies remain beneficial when the cost of infection exceeds the cost of avoiding parasites (Poulin et al. 1999), but organisms should benefit from the ability to modulate their risk avoidance behaviour in relation to the actual threat level (Behringer et al. 2018). Blue mussels clearly show such an ability to adapt their risk behaviour depending on previous experience with trematodes and upregulate their avoidance behaviour when parasite pressure is high. It is unclear, however, if *M. edulis* will ‘unlearn’ the fear of parasites again after prolonged periods of no contact with infective parasite stages. Infections of *H. elongata* in their first intermediate snail host and the release of cercariae into the environment typically peak during the warm summer months and recede in winter (Thieltges & Rick 2006). It is therefore feasible that mussels experience cyclical avoidance learning phases each summer, followed by ‘forgetting’ periods during the winter.

Moreover, recent studies have suggested that fish (sea trout *Salmo trutta trutta*) can change their defence strategies against trematodes (*Diplostomum pseudospathaceum*) over time and shift from avoidance learning to immunological defences after repeated exposure (Klemme & Karvonen 2016). The present study can only present a snapshot in the temporal dynamics of parasite trait responses, which encompasses pre-contact, contact and post-contact phases of host-parasite interaction (Daversa et al. 2021). It therefore remains to be tested, if bivalves can show a similar plasticity in their defence against parasites, beyond avoidance learning. Even more than prey, who often have several encounters with predators and learn to adjust their behaviour accordingly (Laundré et al. 2010), blue mussels will have regular contact with trematode cercariae in their environment that will elicit defensive trait responses. It has been suggested that hosts generally have a broader and more diverse toolkit for defending against parasites due to this continuous interaction than prey have for defending against predators (Daversa et al. 2021), which should be true for the *Mytilus*-trematode system as well.

As an ecosystem engineer that provides relevant ecological functions in coastal habitats, the parasite-mediated lower filtration rates of blue mussels are expected to influence the energy flow in coastal systems (Selbach & Mouritsen 2020). Based on the small-scale experiments in our study, it remains difficult to quantify how and to what extent the observed learning patterns of avoidance behaviour will affect blue mussels at the community and population level, and their overall ecosystem engineering potential, in particular the ability to remove organic matter from the water column. Overall, the learning patterns of mussels in regions with a high abundance of infected snails and high numbers of cercariae in the water, e.g., in shallow waters with high shorebird and periwinkle abundances (Poulin & Mouritsen 2003), should be expected to be different from mussel populations facing only low parasite pressure, e.g., in deeper waters were periwinkles are less common. Based on our findings, it could be expected that mussels regularly facing trematode larvae in the environment become more risk averse than conspecifics in deeper waters. After regular parasite encounters, these populations could show different risk behaviours. Furthermore, within a community of mussels, not all individuals would be expected to show similar levels of avoidance behaviour based on varying experience levels and differences in personality, with less risk-averse mussels being more prone to infections (Barber & Dingemanse 2010). This could be a contributing factor to the often uneven and patchy distribution of trematode infections in mussel populations (Galaktionov et al. 2015).

Naturally, *H. elongata* is not the only threat blue mussels are regularly facing in their aquatic environment. Another trematode species, *Renicola roscovita*, also uses *Mytilus* as an intermediate host but encysts in gills and palps, where it directly impairs the mussels’ filtration ability (Stier et al. 2015). In intertidal mussel beds, both trematode species occur in sympatry and often infect the same mussels (Thieltges & Rick 2006). Since cercariae of both trematodes infect their bivalve host via similar infection pathways, learning to defend against one species might help to avoid another, thereby indirectly shaping parasite transmission dynamics of multiple species. Besides the risks of parasitism, a wide range of predators (birds, crabs, sea stars) prey on *Mytilus edulis* in the intertidal zone. Bivalves have been shown to be capable of sensing and responding to risk cues from predators or alarm cues from conspecifics that face predation pressure (Reimer & Tedengren 1997; Kobak & Ryńska 2014; Dzierżyńska-Białończyk et al. 2019). Accordingly, future research should investigate potential additive or interactive fear effects of the simultaneous presence of parasites and predators in the context of avoidance learning (Daversa et al. 2021).

Overall, non-consumptive and non-lethal effects of predators and parasites can have far-reaching cascading impacts on organisms in an ecosystem, and can even exceed the consumptive effects of these organisms. As we are increasingly uncovering how fear can shape ecosystems, the dynamic learning processes of avoidance behaviour that are likely playing an important role in these interspecific interactions and their ecological impact need to be studied in this context. Blue mussels and their trematode parasites provide an attractive and ecologically important model system to further explore these fundamental questions.

## Ethics

All institutional and national regulations for the care and use of animals were followed.

## Data availability

The datasets supporting this article have been uploaded as part of the electronic supplementary material.

## Author contributions

CS and KNM conceived the study, designed the study and wrote the manuscript. LM, CS and KNM performed the experiments; KNM analysed the data. All authors gave final approval for publication and agree to be held accountable for the work performed therein.

## Competing interests

We declare we have no competing interests.

## Funding

This work received funding from the European Union’s Horizon 2020 Research and Innovation Programme under the Marie Skłodowska-Curie grant agreement No. 839635 TPOINT (CS).

## Acknowledgements

We are grateful to the Danish Shellfish Centre, Mors, Denmark for providing us with blue mussels for the experiment. We thank the Parasite Discussion Club at the IGB Berlin for a constructive discussion of our results, and Jessica Schwelm for feedback on an earlier draft of the manuscript.

## Notes

### Competing Interest Statement

The authors have declared no competing interest.

## References

Adolphs R. 2013 The biology of fear. Curr. Biol. 23, R79–R93. (doi:10.1016/j.cub.2012.11.055)

Bakhmet I, Nikolaev K, Levakin I. 2017 Effect of infection with metacercariae of Himasthla elongata (Trematoda: Echinostomatidae) on cardiac activity and growth rate in blue mussels (Mytilus edulis) in situ. J. Sea Res. 123, 51–54. (doi:10.1016/j.seares.2017.03.012)

Barber I, Dingemanse NJ. 2010 Parasitism and the evolutionary ecology of animal personality. Phil. Trans. R. Soc. B 365, 4077–4088. (doi:10.1098/rstb.2010.0182)

Behringer DC, Karvonen A, Bojko J. 2018 Parasite avoidance behaviours in aquatic environments. Philos. Trans. R. Soc. B Biol. Sci. 373, 20170202. (doi:10.1098/rstb.2017.0202)

Bertness DM, 2007. Atlantic shorelines: natural history and ecology. Princeton University Press, Oxfordshire.

Brown JS, Laundre JW, Gurung M. 1999 The ecology of fear: optimal foraging, game theory, and trophic interactions. J. Mammal. 80, 385–399. (doi:10.2307/1383287)

Buck BH, Thieltges DW, Walter U, Nehls G, Rosenthal H. 2005 Inshore-offshore comparison of parasite infestation in Mytilus edulis: implications for open ocean aquaculture. J. Appl. Ichthyol. 21, 107–113. (doi:10.1111/j.1439-0426.2004.00638.x)

Buck JC, Weinstein SB, Young HS. 2018 Ecological and evolutionary consequences of parasite avoidance. Trends Ecol. Evol. 33, 619–632. (doi:10.1016/j.tree.2018.05.001)

Commito JA, Como S, Grupe BM, Dow WE. 2008 Species diversity in the soft-bottom intertidal zone: biogenic structure, sediment, and macrofauna across mussel bed spatial scales. J. Exp. Mar. Bio. Ecol. 366, 70–81. (doi:10.1016/j.jembe.2008.07.010)

Creel S, Christianson D. 2008 Relationships between direct predation and risk effects. Trends Ecol. Evol. 23, 194–201. (doi:10.1016/j.tree.2007.12.004)

Daversa DR, Hechinger RF, Madin E, Fenton A, Dell AI, Ritchie EG, Rohr J, Rudolf VHW, Lafferty KD. 2021 Broadening the ecology of fear: non-lethal effects arise from diverse responses to predation and parasitism. Proc. R. Soc. B Biol. Sci. 288, 20202966. (doi:10.1098/rspb.2020.2966)

Dzierżyńska-Białończyk A, Jermacz Ł, Zielska J, Kobak J. 2019 What scares a mussel? Changes in valve movement pattern as an immediate response of a byssate bivalve to biotic factors. Hydrobiologia 841, 65–77. (doi:10.1007/s10750-019-04007-0)

Ford AT, Goheen JR, Otieno TO, Bidner L, Isbell LA, Palmer TM, Ward D, Woodroffe R, Pringle RM. 2014 Large carnivores make savanna tree communities less thorny. Science. 346, 346–349. (doi:10.1126/science.1252753)

Galaktionov K V. et al. 2015 Factors influencing the distribution of trematode larvae in blue mussels Mytilus edulis in the North Atlantic and Arctic Oceans. Mar. Biol. 162, 193–206. (doi:10.1007/s00227-014-2586-4)

James CT, Noyes KJ, Stumbo AD, Wisenden BD, Goater CP. 2008 Cost of exposure to trematode cercariae and learned recognition and avoidance of parasitism risk by fathead minnows Pimephales promelas. J. Fish Biol. 73, 2238–2248. (doi:10.1111/j.1095-8649.2008.02052.x)

Klemme I, Karvonen A. 2016 Learned parasite avoidance is driven by host personality and resistance to infection in a fish–trematode interaction. Proc. R. Soc. B Biol. Sci. 283, 20161148. (doi:10.1098/rspb.2016.1148)

Kobak J, Ryńska A. 2014 Environmental factors affecting behavioural responses of an invasive bivalve to conspecific alarm cues. Anim. Behav. 96, 177–186. (doi:10.1016/j.anbehav.2014.08.014)

Koprivnikar J, Penalva L. 2015 Lesser of two evils? Foraging choices in response to threats of predation and parasitism. PLoS One 10, e0116569. (doi:10.1371/journal.pone.0116569)

Larsen MH, Høeg JT, Mouritsen KN. 2013 Influence of infection by Sacculina carcini (Cirripedia, Rhizocephala) on consumption rate and prey size selection in the shore crab Carcinus maenas. J. Exp. Mar. Bio. Ecol. 446, 209–215. (doi:10.1016/j.jembe.2013.05.029)

Larsson J, Lind EE, Corell H, Grahn M, Smolarz K, Lönn M. 2017 Regional genetic differentiation in the blue mussel from the Baltic Sea area. Estuar. Coast. Shelf Sci. 195, 98–109. (doi:10.1016/j.ecss.2016.06.016)

Lauckner G. 1983. Diseases of mollusca: bivalvia. In: Diseases of Marine Animals, Vol. II, ed. O. Kinne (Hamburg: Biologische Anstalt Helgoland), 477–961.

Laundré JW, Hernandez L, Ripple WJ. 2010 The landscape of fear: ecological implications of being afraid. Open Ecol. J. 3, 1–7. (doi:10.2174/1874213001003030001)

Marino JA, Werner EE. 2013 Synergistic effects of predators and trematode parasites on larval green frog (Rana clamitans) survival. Ecology 94, 2697–2708. (doi:10.1890/13-0396.1)

Marras S, Domenici P. 2013 Schooling fish under attack are not all equal: some lead, others follow. PLoS One 8, e65784. (doi:10.1371/journal.pone.0065784)

Mascaró M, Seed R. 2001 Choice of prey size and species in Carcinus maenas (L.) feeding on four bivalves of contrasting shell morphology. Hydrobiologia 449, 159–170. (doi:10.1023/A:1017569809818)

Mouritsen KN, Dalsgaard NP, Flensburg SB, Madsen JC, Selbach C. 2021 Fear of parasitism affects the functional role of ecosystem engineers. bioRxiv (doi:10.1101/2021.08.27.457894)

Mouritsen KN. 2017 Periwinkle regulation: parasitism and epibiosis are linked. Mar. Ecol. Prog. Ser. 579, 227–231. (doi:10.3354/meps12277)

Peacor SD, Werner EE. 2001 The contribution of trait-mediated indirect effects to the net effects of a predator. Proc. Natl. Acad. Sci. U. S. A. 98, 3904–3908. (doi:10.1073/pnas.071061998)

Poulin R, Marcogliese DJ, McLaughlin JD. 1999 Skin-penetrating parasites and the release of alarm substances in juvenile rainbow trout. J. Fish Biol. 55, 47–53. (doi:10.1006/jfbi.1999.0970)

Poulin R, Mouritsen K. 2003 Large-scale determinants of trematode infections in intertidal gastropods. Mar. Ecol. Prog. Ser. 254, 187–198. (doi:10.3354/meps254187)

Preisser EL, Bolnick DI, Benard MF. 2005 Scared to death? The effects of intimidation and consumption in predator-prey interactions. Ecology 86, 501–509. (doi:10.1890/04-0719)

Ragnarsson SÁ, Raffaelli D. 1999 Effects of the mussel Mytilus edulis L. on the invertebrate fauna of sediments. J. Exp. Mar. Bio. Ecol. 241, 31–43. (doi:10.1016/S0022-0981(99)00063-5)

Reimer O, Tedengren M. 1997 Predator-induced changes in byssal attachment, aggregation and migration in the blue mussel, Mytilus edulis. Mar. Freshw. Behav. Physiol. 30, 251–266. (doi:10.1080/10236249709379029)

Riemann B, Simonsen P, Stensgaard L. 1989 The carbon and chlorophyll content of phytoplankton from various nutrient regimes. J. Plankton Res. 11, 1037–1045. (doi:10.1093/plankt/11.5.1037)

Ripple WJ, Beschta RL. 2004 Wolves and the ecology of fear: can predation risk structure ecosystems? Bioscience 54, 755. (doi:10.1641/0006-3568(2004)054[0755:wateof]2.0.co;2)

Rohr JR, Swan A, Raffel TR, Hudson PJ. 2009 Parasites, info-disruption, and the ecology of fear. Oecologia 159, 447–454. (doi:10.1007/s00442-008-1208-6)

Selbach C, Mouritsen KN. 2020 Mussel shutdown: does the fear of trematodes regulate the functioning of filter feeders in coastal ecosystems? Front. Ecol. Evol. 8. (doi:10.3389/fevo.2020.569319)

Silva BA, Gross CT, Gräff J. 2016 The neural circuits of innate fear: detection, integration, action, and memorization. Learn. Mem. 23, 544–555. (doi:10.1101/lm.042812.116)

Stier T, Drent J, Thieltges D. 2015 Trematode infections reduce clearance rates and condition in blue mussels Mytilus edulis. Mar. Ecol. Prog. Ser. 529, 137–144. (doi:10.3354/meps11250)

Suraci JP, Clinchy M, Dill LM, Roberts D, Zanette LY. 2016 Fear of large carnivores causes a trophic cascade. Nat. Commun. 7, 10698. (doi:10.1038/ncomms10698)

Thieltges DW, Rick J. 2006 Effect of temperature on emergence, survival and infectivity of cercariae of the marine trematode Renicola roscovita (Digenea: Renicolidae). Dis. Aquat. Organ. 73, 63–8. (doi:10.3354/dao073063)

Weinstein SB, Moura CW, Mendez JF, Lafferty KD. 2018 Fear of feces? Tradeoffs between disease risk and foraging drive animal activity around raccoon latrines. Oikos 127, 927– 934. (doi:10.1111/oik.04866)

Werding B. 1969 Morphologie, Entwicklung und Ökologie digener Trematoden-Larven der Strandschnecke Littorina littorea. Mar. Biol. 3, 306–333. (doi:10.1007/BF00698861)

Zanette LY, Clinchy M. 2019 Ecology of fear. Curr. Biol. 29, R309–R313. (doi:10.1016/j.cub.2019.02.042)

